# Quantitative characterisation of low abundant yeast mitochondrial proteins reveals compensation for haplo-insufficiency in different environments

**DOI:** 10.1101/2021.07.09.451775

**Authors:** Alkisti Manousaki, James Bagnall, David Spiller, Laura Natalia Balarezo-Cisneros, Michael White, Daniela Delneri

## Abstract

Quantification of low abundant membrane-binding proteins such as transcriptional factors and chaperones has been proved difficult even with the most sophisticated analytical technologies. Here we exploit and optimise the non-invasive Fluorescence Correlation Spectroscopy (FCS) for quantitation of low abundance proteins and as proof of principle we choose two interacting proteins involved in fission of mitochondria in yeast, Fis1p and Mdv1p. In *Saccharomyces cerevisiae* the recruitment of Fis1p and Mdv1p to mitochondria is essential for the scission of the organelles and the retention of functional mitochondrial structures in the cell. We used FCS in single, GFP-labelled live yeast cells to quantify the protein abundance in homozygote and heterozygote cells, and to investigate the impact of the environments on protein copy number, bound/unbound protein state and mobility kinetics. Both proteins were observed to localise predominantly at mitochondrial structures with the Mdv1p bound state increasing significantly in a strictly respiratory environment. Moreover, a compensatory mechanism which controls Fis1p abundance upon deletion of one allele was observed in Fis1p but not in Mdv1p, suggesting differential regulation of Fis1p and Mdv1p protein expression.

## Introduction

Biological systems are complex and require the dynamic and coordinated function of various proteins that may operate differently within the same cell or a population of cells. Identical cells of the same population have been shown to respond heterogeneously to a variety of inter- and intra-cellular signals within seconds or minutes [1–3]. The variability in cellular responses can include changes in gene expression, protein transcription and translation, interactions between proteins and molecule kinetics [3]. To better understand the dynamic nature of biological processes, it is essential to precisely define the concentration and mobility rates of the proteins involved in real time and at the single-cell level. Most of the available technologies allow precise quantification of inducible or inherently abundant proteins for measurements performed on cell populations or synchronised cells [4–7]. In contrast, proteins of low abundance and endogenous expression often miss detection or are defined as noise, conferring the major bottleneck in protein quantification to date [8–10].

The use of high-throughput microscopy technologies coupled with advanced software tools and mathematical modelling offers a new potential for defining the dynamics of low-abundant proteins in live, single cells. Fluorescent Correlation Spectroscopy (FCS) is a non-invasive microscopy technique which can achieve absolute quantification of fluorescent particles as they diffuse at low numbers within the cells by measurements taken at multiple time points and under different conditions [11–13]. In its principle, FCS investigates the stochasticity in protein expression as this is exhibited by fluctuations in fluorescence intensity that emerge from the transition of labelled molecules through a diffraction limited focal volume of light [14, 15]. Successively, the fluorescent signal emitted from the illuminated area is recorded over time in orders of sub-seconds to minutes and analysed via autocorrelation functions into numbers of mobility and molecules per cell [12]. Additional FCS analysis on the diffusion rate of the labelled particles can provide information about the size of the molecules or their localisation state; large or membrane-bound molecules diffuse slower than those that are smaller in size or move freely [16, 17].

Much of the progress achieved in quantitative proteomics over the years results from microscopy-based research conducted in yeast, with *Saccharomyces cerevisiae* having a well-defined numerical identity and localization pattern for over 5000 out of its 6.600 proteins [18–29]. The construction of the yeast GFP fusion library conferred a significant advantage to this direction as it covers approximately 70% of the yeast proteome [6]. Findings have shown that yeast proteins range between 3 to 7.5 × 10^5^ copies per cell, with 67% of them measured between 1000 and 10000 molecules with a median abundance of 2622 copies per cell [29]. Proteins present in the cell at 866 or fewer copies are defined as molecules of low abundance in contrast to those of high abundance which are found at 1.4 x 10^5^ copies or more [29]. Approximately 1000 of the yeast proteins comprise the mitochondrial proteome, which involves less abundant, nuclear-encoded (> 90%) protein molecules of diverse functions [29]. Among these mitochondrial proteins are the Fis1p and Mdv1p fission components, which are less studied in terms of their spatial and temporal dynamics mainly due to their low expression.

The Fis1p and Mdv1p yeast proteins regulate the scission of mitochondria through direct interaction with the Dnm1p dynamin-related GTPase [30–32]. Dnm1p is recruited to mitochondria by Fis1p which anchors to the outer mitochondrial membrane through a single transmembrane region at its C-terminus [30, 33–35]. The N-terminal of Fis1p remains in the cytosol and recruits the adaptor protein Mdv1p or its paralog Caf4p when Mdv1p is not present [32, 36, 37]. At the early stages of fission and in the absence of Dnm1p, both Fis1p and Mdv1p are evenly distributed on the outer mitochondrial surface [31]. However, during fission Mdv1p co-assembles with Dnm1p into punctate structures at sites of future mitochondrial constriction [31]. The recruitment of Fis1p to mitochondria is essential for the distribution and function of the Dnm1p::Mdv1p fission complexes, but remains independent from both Dnm1p and Mdv1p [30]. Apart from its role in mitochondrial fission, membrane-bound forms of Fis1p also tether damaged or misfolded proteins to mitochondria of the mother cells [38], mediate their apoptotic fragmentation in response to ethanol [39, 40] and promote cell death in aging cells [41]. Furthermore, Fis1p is an ambiguous protein [42, 43], which means that it can be recognised by more than one cellular departments, and has a portion of cytosolic copies localised on peroxisomes to regulate their abundance [44, 45]. Similarly to Fis1p, the majority of Mdv1p is associated with mitochondria [31] whereas a portion of the protein is involved in the regulation of peroxisomal fission [45]. Notably, no homologs of Mdv1p are found in other organisms than fungi.

Several attempts have been made over the last years to provide quantitative information on the abundance and composition of the Fis1p and Mdv1p proteins involved in mitochondrial division [6, 7, 25, 46]. Interestingly though, several studies in the literature provide different localisation patterns and copy numbers for these fission components (SGD database, [46]). In this study, we used and optimised the state-of-the-art fluorescence spectroscopy technique, FCS, to precisely quantify the native expression of *FIS1* and *MDV1* at the protein level as this is regulated by the endogenous promoters of the alleles. We were able to determine quantitatively the intracellular and mitochondrial-related protein abundance, the physical state of the proteins (bound and unbound forms), the heterogeneity in protein copy numbers within the populations of homozygote and heterozygote yeast cells. Interestingly, we showed compensation in levels of protein abundance upon deletion of one copy of the *FIS1* gene, and a distinct behaviour of the proteins in response to different environmental conditions.

## Results

### Generation of the yeast GFP fusion strain collection

The use of the FCS microscopy method requires the utilisation of fluorescent fusion proteins. Beside strain *Sc* FIS1^GFP^ *MAT*a and *Sc* MDV1^GFP^ *MAT*a that were obtained from the GFP-tagged collection, the six additional strains were constructed expressing the C-terminally GFP-tagged Fis1p and Mdv1p mitochondrial fission proteins under the control of their own native promoters (Table 1). Correct chromosomal insertion of the protein fluorophores was confirmed by PCR and Sanger sequencing (Table S1). Strains were verified for their ploidy by *MAT* locus PCR and FACS analysis (Table S1, Fig. S1). Furthermore, we checked whether the tag interfere with the expression of the labelled Fis1p and Mdv1p molecules. No difference was observed on the expression of the total Fis1p-GFP and Mdv1p-GFP in haploid *S. cerevisiae* cells growing in glucose when compared to the non-tagged Fis1p and Mdv1p proteins, respectively (Fig. S2).

**Table 1.**
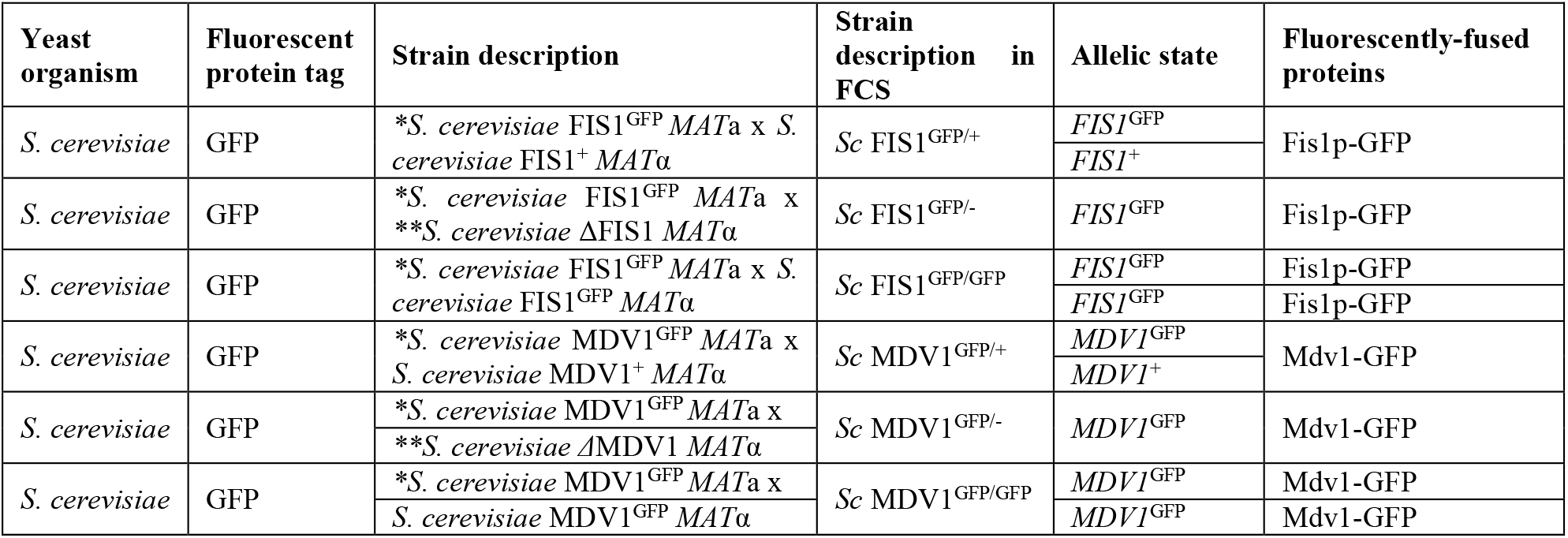
Yeast library of GFP fusion strains. The table contains all strains constructed and/or manipulated in this study along with information regarding the yeast background, type of fluorophore, strain genotype, genes fused to fluorophores and fusion proteins expressed. * Indicates strains purchased from the GFP-tagged collection (Life Technologies, UK, Clone ID: YILO65C and YJL112W), ** indicates mutant strains from Yeast Deletion Collection available in the Delneri lab, *Sc* represent an *S. cerevisiae* strain background.

### Quantification of Fis1p and Mdv1p heterogeneity in single cells of *S. cerevisiae* populations

Protein dynamics can be heterogeneous in terms of protein expression, molecule concentration and mobility among identical cells of the same population [1–3]. To quantitatively determine the inherent heterogeneity of Fis1p and Mdv1p protein expression, we measured the intracellular abundance of endogenously expressed Fis1p-GFP and Mdv1p-GFP in live, single cells of *S. cerevisiae* populations using FCS (Fig. 1).

**Figure 1.**
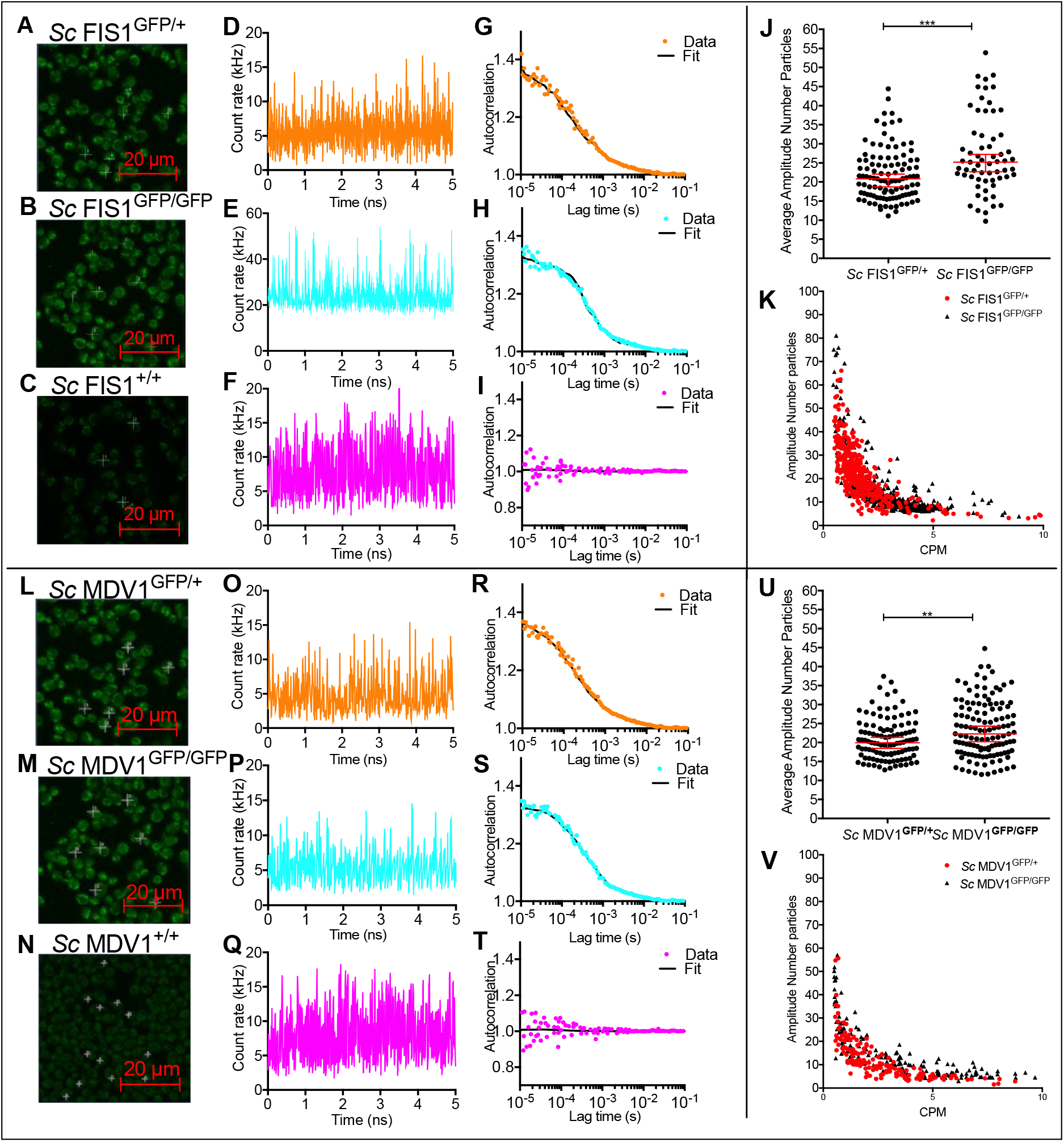
Quantification of Fis1p-GFP and Mdv1p-GFP heterogeneity in living yeast cells. Cells of *Sc* FIS1^GFP/+^, *Sc* FIS1^GFP/GFP^ and *Sc* FIS1^+/+^ were imaged under laser excitation at 488 nm (A-C). Fluorescence fluctuation traces (fluorescent count in kHz) were obtained over a 5s period by FCS measurements performed on mitochondrial locations of exponentially growing *Sc* FIS1^GFP/+^ (n = 107) (D), *Sc* FIS1^GFP/GFP^ (n = 63) (E) and *Sc* FIS1^+/+^ (n = 84) (F) cells. Correlation analysis showed positive autocorrelation of Fis1p-GFP in *Sc* FIS1^GFP/+^ (G) and *Sc* FIS1^GFP/GFP^ (H) cells and gave no amplitude for the GFP-free *Sc* FIS1^+/+^ control cells (I). Scatter plots of mitochondrial Fis1p-GFP concentrations calculated by FCS measurements taken in individual *Sc* FIS1^GFP/+^ (n = 107) and *Sc* FIS1^GFP/GFP^ (n = 63) cells (J). The CPM protein quantification values are negatively and linearly correlated with the Amplitude for both cell types (K), indicating no fluorescence artefacts (*Sc* FIS1^GFP/+^, R^2^ = 0.41 and *Sc* FIS1^GFP/GFP^, R^2^ = 0.42). Cells of *Sc* MDV1^GFP/+^, *Sc* MDV1^GFP/GFP^ and *Sc* MDV1^+/+^ were imaged under laser excitation at 488 nm (L-N). Fluorescence fluctuation traces (fluorescent count in kHz) were obtained over a 5 second period by FCS measurements performed on mitochondrial locations of exponentially growing *Sc* MDV1^GFP/+^ (n = 108) (O), *Sc* MDV1^GFP/GFP^ (n = 124) (P) and *Sc* MDV1^+/+^ (n = 84) (Q) cells. Correlation analysis showed positive autocorrelation of Mdv1p-GFP in *Sc* MDV1^GFP/+^ (R) and *Sc* MDV1^GFP/GFP^ (S) cells and gave no amplitude for the GFP-free *Sc* MDV1^+/+^ control cells (T). Scatter plots of mitochondrial Mdv1p-GFP concentrations calculated by FCS measurements taken in individual Sc MDV1^GFP/+^ (n = 108) and Sc MDV1^GFP/GFP^ (n = 124) cells (U). The CPM protein quantification values are negatively and linearly correlated with the Amplitude for both cell types (V), indicating no fluorescence artefacts (*Sc* MDV1^GFP/+^, R^2^ = 0.45 and *Sc* MDV1^GFP/GFP^, R^2^ = 0.52). Scale bar is at 20 μm. The upper and lower 95% confidence intervals are shown as error bars that extend above and below the top of the median bar. *** represents significance at the 0.001 level and ** at the 0.01 level for the Mann Whitney two-tailed t-test.

We determined the level of sensitivity in FCS measurements by measuring the variation in low Fis1p-GFP protein copies in cells growing naturally in rich fermentable carbon sources. Two diploid strains, one having both *FIS1* alleles tagged to GFP (*Sc* FIS1^GFP/GFP^) and one having only one allele tagged to GFP (*Sc* FIS1^GFP/+^), were imaged under laser excitation at 488 nm and compared to the negative control strain (*Sc* FIS1^+/+^) carrying no GFP markers (Fig. 1, A-C). A mitochondrial localisation of Fis1p-GFP inside the cytoplasm was determined for exponentially growing cells of both *Sc* FIS1^GFP/+^ and *Sc* FIS1^GFP/GFP^ populations (Fig. 1, A and B). To quantify the level of signal emission, we calculated the fluctuations in GFP fluorescence as a function of time from measurements taken on mitochondrial locations and we generated individual measurements of counts per molecule (CPM) for each set of strains (Fig. 1, D-F). The CPM of the GFP-fused Fis1p in the *Sc* FIS1^GFP/GFP^ cells (n = 63) was four-fold higher (mean molecular brightness of 20; SD±1.1) than in *Sc* FIS1^GFP/+^ cells (n = 107; mean molecular brightness of 5; SD±1.2) (Fig. 1, D and E). In contrast, the fluorescence fluctuation trace generated for the non-labelled *Sc* FIS1^+/+^ cells (n = 93) showed no characteristic peak for the GFP signal (Fig. 1, F).

To determine whether the difference observed in GFP fluorescence between the *Sc* FIS1^GFP/GFP^ and *Sc* FIS1^GFP/+^ cells corresponds to higher levels of protein abundance, we generated the FCS autocorrelation curves (Fig. 1, G and H). Firstly, we found the amplitude starting point of the autocorrelation curve for the *Sc* FIS1^GFP/+^ cells (n = 107) and for the *Sc* FIS1^GFP/GFP^ cells (n = 63) to be 1.37 (Fig. 1, G) and 1.32 (Fig. 1, H), respectively, suggesting a lower degree of normalised variance for the latter strain. Secondly, we determined the levels of cellular autofluorescence by comparing the fluorescence fluctuation traces obtained for the GFP labelled cells to the non-labelled strain *Sc* FIS1^+/+^ (Fig. 1, I). The autocorrelation curve for the *Sc* FIS1^+/+^ cells showed no characteristic amplitude decay over the lag time, confirming the absence of green signal (Fig. 1, I). Finally, in order to quantify the intercellular heterogeneity of Fis1p in each set of strains, we calculated the total number of GFP-fused Fis1p diffusing through the confocal volume (CV) (Fig. 1, J). The median number of Fis1p-GFP inside the confocal volume was measured to be 25.19 (95% CI: 22.66 - 27.22) for the *Sc* FIS1^GFP/GFP^ cells (n = 63) and 20.85 (95% CI: 18.74 - 22.04) for the *Sc* FIS1^GFP/+^ cells (n = 107). This data suggests that the strains with both alleles tagged have as expected a higher number of measured proteins compared to the strains with only one allele tagged. The median molecule numbers of each cell population were compared using a non-parametric Mann-Whitney two-tailed t-test, and the average number of GFP-fused Fis1p was indeed significantly higher (p = 0.0002) in the *Sc* FIS1^GFP/GFP^ cells when compared to *Sc* FIS1^GFP/+^ cells (Fig. 1, J). To ensure that this comparison was not affected by some *a priori* differential relationship of the CPM and the amplitude FCS data, the obtained R^2^ coefficients were compared in a fitted linear regression model (Fig. 1, K). No differences were observed between the *Sc* FIS1^GFP/GFP^ (R^2^: 0.42) and *Sc* FIS1^GFP/+^ (R^2^: 0.41) cells.

The same experimental strategy was used for Mdv1p (Fig. 1, L-V) and similar results were obtained. In this case we found that the median number of Mdv1p-GFP molecules associated with mitochondria is 22.29 (95% CI: 20.22 - 24.32) and 19.93 (95% CI: 18.48 - 21.35) in the *Sc* MDV1^GFP/GFP^ (n = 124) and *Sc* MDV1^GFP/+^ (n = 108) cells, respectively. This difference is also significant (Man Whitney two-tailed t-test, p = 0.0072) with no differential R^2^ coefficient for CPM and the amplitude FCS data between *Sc* MDV1^GFP/GFP^ (R^2^: 0.52) and *Sc* MDV1^GFP/+^ (R^2^: 0.45).

### Compensatory mechanisms are present in heterozygote mutants for Fis1p in fermentative and respiratory conditions

Having defined the level of Fis1p-GFP and Mdv1p-GFP heterogeneity within the population, we next explored the effect of gene copy number on the intracellular abundance of the labelled proteins in single live cells. We compared the levels of Fis1p-GFP and Mdv1p-GFP proteins in different GFP-tagged *S. cerevisiae* strains having the second, non-labelled allele deleted (*i.e*. strains *Sc* FIS1^GFP/-^ and *Sc* MDV1^GFP/-^) with the measurements obtained for the strains *Sc* FIS1^GFP/+^ and *Sc* MDV1^GFP/+^, where both alleles were present (Fig. 2).

**Figure 2.**
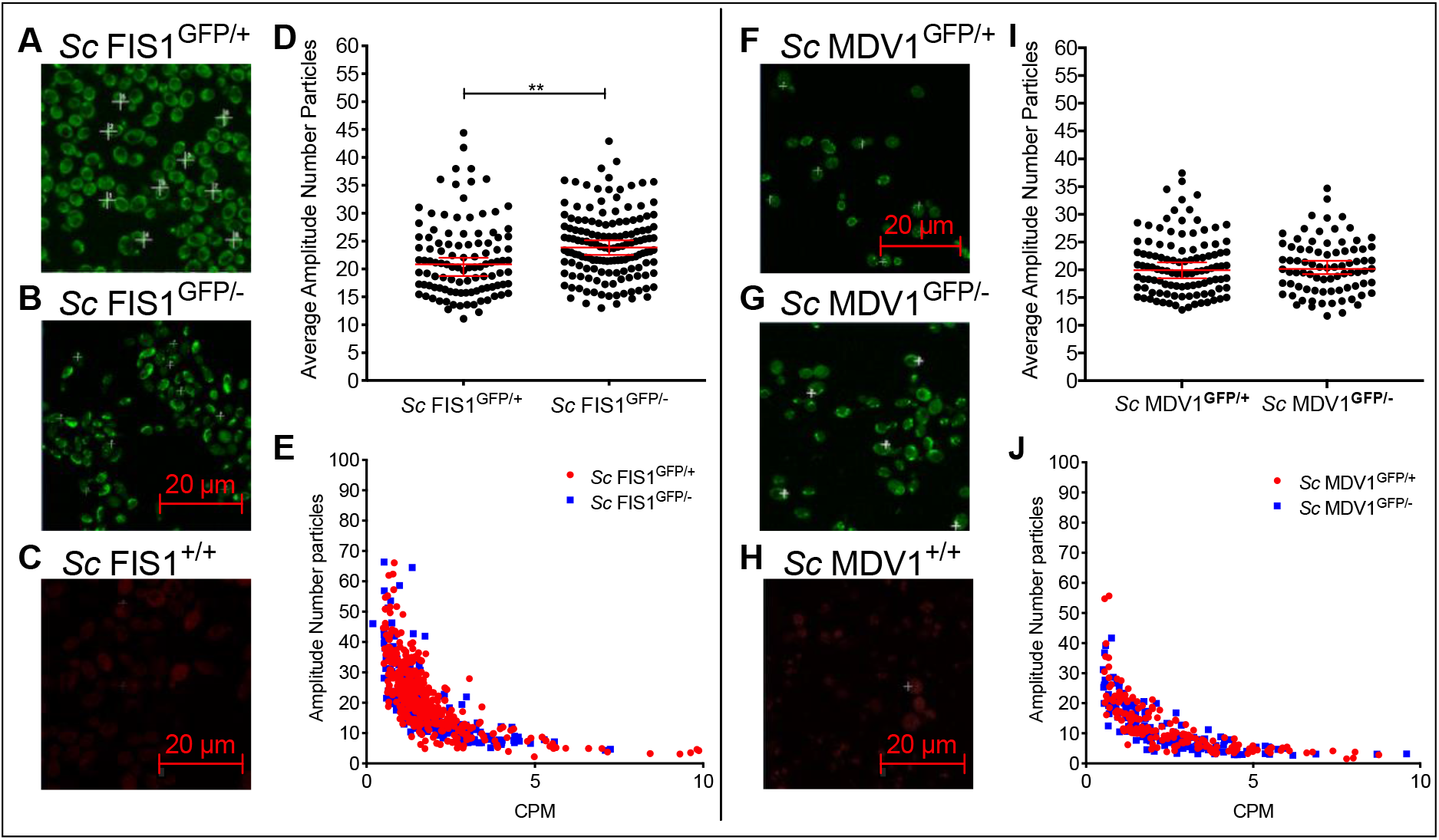
Detection of increased mitochondrial-associated Fis1p-GFP and Mdv1p-GFP in heterozygote yeasts by FCS. Example images of *Sc* FIS1^GFP/+^ (A), *Sc* FIS1^GFP/-^ (B) and *Sc* FIS1^+/+^ (C) show yeast cells of an OD_600_ of 0.5 in SD media containing 2% glucose imaged under simultaneous laser excitation at 488 nm and 561 nm (Scale bar = 20 μm). Panel (D) presents the scatter plots showing the heterogenous expression of Fis1p-GFP measured in single cells of *Sc* FIS1^GFP/+^ (n = 107) and *Sc* FIS1^GFP/-^ (n = 145) cells. In panel (E), the correlation between the CPM and amplitude values reveals no technical artefacts for FCS measurements taken in both cell lines (*Sc* FIS1^GFP/+^, R^2^ = 0.41 and *Sc* FIS1^GFP/-^, R^2^ = 0.43). Images (F-H) show exponentially growing *Sc* MDV1^GFP/+^ (F), *Sc* MDV1^GFP/+^ (G), *Sc* MDV1^+/+^ (H) yeast cells in SD media containing 2% glucose and under simultaneous laser excitation at 488 nm and 561 nm (Scale bar = 20 μm). Scatter plot in panel (I) shows the heterogenous levels of Mdv1p-GFP mitochondrial abundance measured in single cells of *Sc* MDV1^GFP/+^ (n = 108) and *Sc* MDV1^GFP/-^ (n = 81) populations. In panel (J), the correlation of CPM to amplitude values reveals no technical artefacts for measurements taken in all cell types (*Sc* MDV1^GFP/+^, R^2^ = 0.46 and *Sc* MDV1^GFP/-^, R^2^ = 0.49). Error bars show the 95% confidence intervals and the median value. Statistical significance was calculated using Man Whitney two-tailed t-test. ** represents significance at the 0.01 level.

To determine the levels of the GFP signal, we imaged exponentially growing *Sc* FIS1^GFP/+^, *Sc* FIS1^GFP/-^ and *Sc* FIS1^+/+^ cells under simultaneous excitation at the green spectrum using FCS (see Methods) (Fig. 2, A-C). We found the median number of Fis1p-GFP protein copies detected within the confocal volume to be 23.85 (95% CI: 22.53 - 25.15) for the *Sc* FIS1^GFP/-^ cells (n = 145) and 20.85 (95% CI: 18.74 - 22.04) for the *Sc* FIS1^GFP/+^ cells (n = 107) (Fig. 2, D). The median GFP-fused Fis1p was significantly higher (Man Whitney two-tailed t-test, p = 0.0004) in the heterozygote mutant *Sc* FIS1^GFP/-^ than the *Sc* FIS1^GFP/+^ (Fig. 2, D). No differences in the R^2^ coefficients were observed in the two cell types (*Sc* FIS1^GFP/-^ R^2^ = 0.43 and *Sc* FIS1^GFP/+^ R^2^ = 0.41; Fig. 2, E), confirming that the data were not influenced by technical artefacts. This result indicates that the strains lacking one *FIS1* allele have higher protein abundance for the mitochondrial-associated Fis1p compared to the strains having both alleles.

In addition, the yeast Fis1p is known to interact directly with Mdv1p to form the Fis1p::Mdv1p mitochondrial fission protein complex [31, 47, 48]. However, Mdv1p does not show a compensatory trend, with median mitochondrial Mdv1p-GFP of 20.19 (95% CI: 19.20 - 21.62) in *Sc* MDV1^GFP/-^ (n = 81) and 19.93 (95% CI: 18.48 - 21.35) in *Sc* MDV1^GFP/+^ (n = 108), respectively (Fig. 2, H-J).

Fis1p and Mdv1p protein abundance was also measured under strict respiratory conditions, in media containing glycerol as sole carbon source, where the function of mitochondria is essential (Fig. 3). We found the mitochondrial concentrations of Fis1p-GFP to differ significantly between the two cell populations (Man Whitney two-tailed t-test, p = 0.034) with the median copy numbers of the confocal mitochondrial proteins of 22.98 (95% Cl: 20.56 - 25.76) and 20.02 (95% Cl: 18.25 - 23.03) for the *Sc* FIS1^GFP/-^ and *Sc* FIS1^GFP/+^ strains, respectively (Fig. 3, D). Collectively, the protein quantification data generated for measurements taken on mitochondrial locations of *S. cerevisiae* cells growing on fermentable (glucose) and respiratory (glycerol) carbon sources, indicate that the mitochondrial Fis1p shows the same compensation trend in levels of protein abundance upon *FIS1* gene deletion. In agreement to the data obtained for the glucose growth, the levels of mitochondrial-associated Mdv1p-GFP did not change (Man Whitney two-tailed t-test; p > 0.05) upon deletion of one *MDV1* allele when quantified in *Sc* MDV1^GFP/+^ and *Sc* MDV1^GFP/-^ strains growing in glycerol (Fig. 3, F-J).

**Figure 3.**
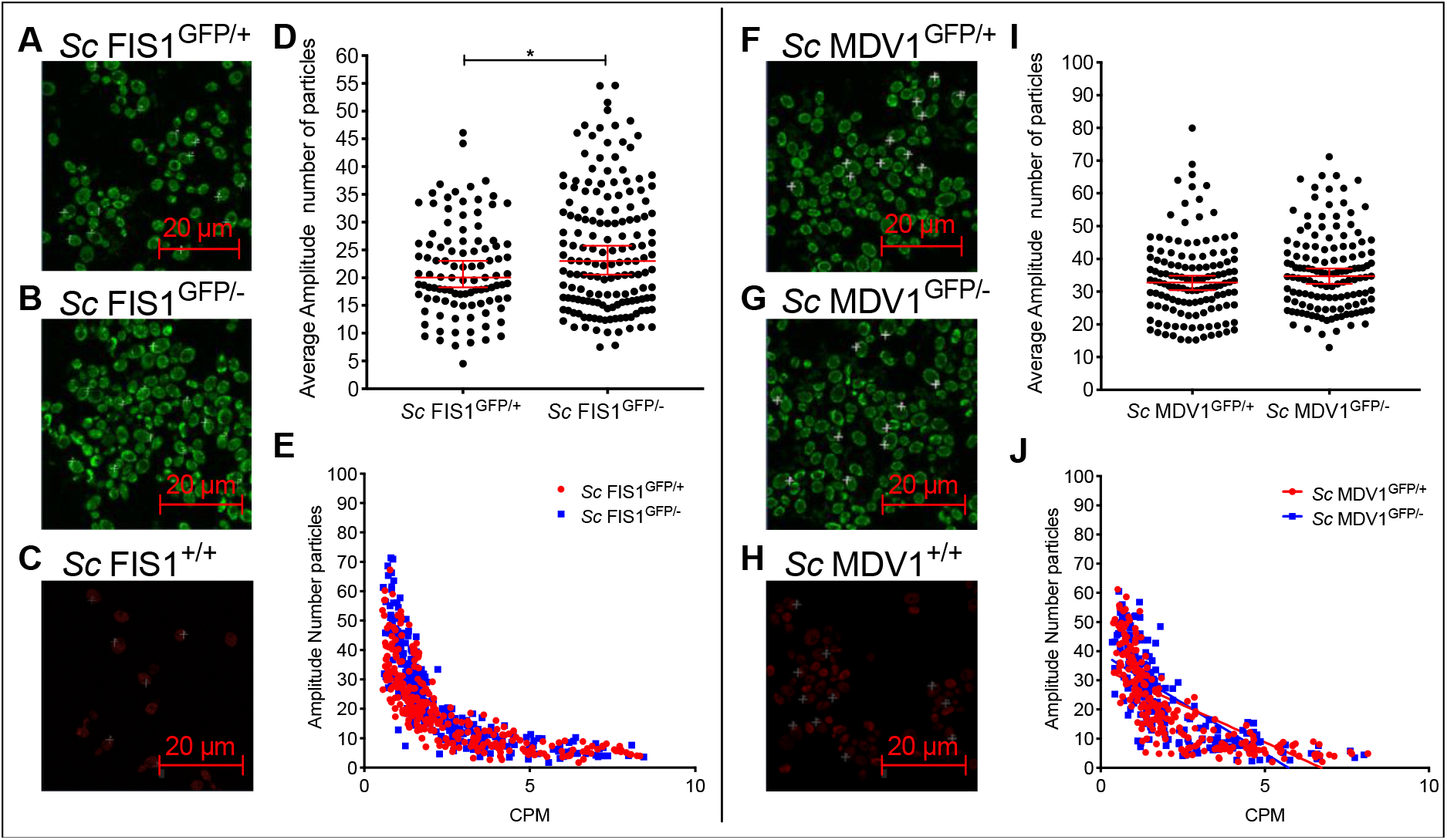
Measurement of Fis1p-GFP and Mdv1p-GFP abundance in yeast cells growing in non-fermentable carbon source. Example images of *Sc* FIS1^GFP/+^ (A), *Sc* FIS1^GFP/+^ (B) and *Sc* FIS1^+/+^ (C) were obtained by imaging cells in the mid-logarithmic phase in media containing glycerol (Scale bar = 20μm). Scatter plots (D) show the heterogeneity in molecule number in confocal volume measured in single cells of *Sc* FIS1^GFP/+^ (n = 103) and *Sc* FIS1^GFP/-^ (n = 158) cell populations. Linear regression graph (E) shows the correlation between the CPM and amplitude values revealing no technical artefacts for FCS measurements taken in both cell lines (*Sc* FIS1^GFP/+^, R^2^ = 0.51 and *Sc* FIS1^GFP/-^, R^2^ = 0.55). Example images of *Sc* MDV1^GFP/+^ (F), *Sc* MDV1^GFP/-^ (G) and *Sc* MDV1^+/+^ (H) were obtained from FCS measurements on exponentially growing cells in media containing glycerol. Scatter plots show the heterogeneity in molecule number in confocal volume measured in single cells of *Sc* MDV1^GFP/+^ (n = 131) and *Sc* MDV1^GFP/-^ (n = 135) cell populations (I). Linear regression graph (J) shows the correlation between the CPM and amplitude values revealing no technical artefacts for FCS measurements taken in both cell lines (*Sc* MDV1^GFP/+^, R^2^ = 0.51 and *Sc* MDV1^GFP/-^, R^2^ = 0.54). Scale bar is at 20μm. Error bars show the 95% confidence intervals and the median value. Statistical significance was calculated using Mann-Whitney two-tailed t-test. * represents significance for p≤0.05.

Interestingly, the concentration of Fis1p-GFP mitochondrial proteins detected in *Sc* FIS1^GFP/+^ growing in glycerol did not differ significantly compared to glucose (Mann Whitney two-tailed t-test, p>0.05), and the same applied for the Fis1p-GFP molecules detected in *Sc* FIS1^GFP/-^ (Mann Whitney two-tailed t-test, p>0.05) (Fig. 4, A). On the other hand, the copy numbers of mitochondrial-associated Mdv1p-GFP in *Sc* MDV1^GFP/+^ and *Sc* MDV1^GFP/-^ were significantly different between the two growth conditions (Man Whitney two-tailed t-test, p<0.0001; Fig. 4, B).

**Figure 4.**
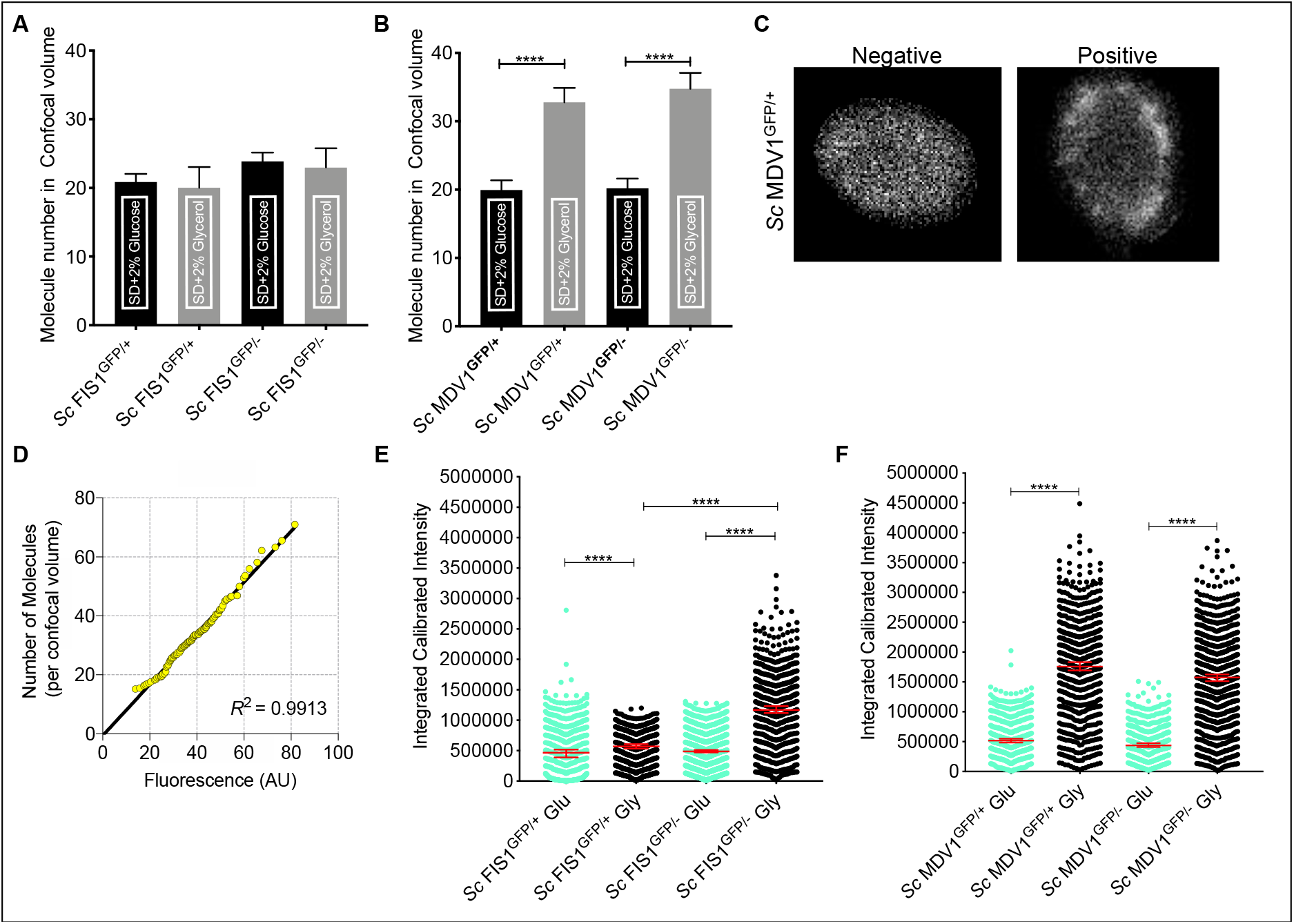
Cellular and mitochondrial-associated Fis1p-GFP and Mdv1p-GFP abundance. Comparison of the mitochondrial-associated Fis1p-GFP (A) and Mdv1p-GFP (B) abundance under different environmental and allelic perturbations. Images (C) showing *Sc* MDV1^GFP/+^ cells: Negative cells demonstrate even distribution of fluorescence across the cell, positive cells show mitochondrial localisation of fluorescence. Mitochondrial localisation was defined after comparison with cells carrying no fluorescence marker and stained mitochondria with Cox4p-DsRed. Example of Quantile-Quantile plot of total and mitochondrial Fis1p-GFP protein distribution across their respective cell populations (D). Comparison of the Integrated calibrated intensities of Fis1p-GFP (E) and Mdv1p-GFP (F) total cell abundance under different environmental and allelic perturbations. Error bars show the 95% confidence intervals and the median value. Statistical significance was calculated using Man Whitney two-tailed t-test. **** represents significance at the 0.0001 level.

To investigate whether the differences observed are specific to the mitochondria location or relate to the total cell protein, we calculated the protein concentration (Fis1p-GFP and Mdv1p-GFP) in the whole cell. Using the acquired confocal microscopy images we measured the fluorescence emitted from the whole cell. First, individual cells were segmented from whole field images in an automated manner. Figure S3 shows 100 randomly selected individual cells for each strain and growth condition from this step. Detection of subcellular organelles was then performed on the entire individual datasets using the Image J Squassh plugin. Object identification parameters were optimised on positive control cells stained for mitochondria with the dye Cox4p-DsRed (see supplementary for parameters conditions used). The same analysis was then applied to all individual cell data sets, which resulted in cells being categorised into positive (subcellular object detected) and negative (even distribution of fluorescence across cell) (Fig. 4, C; Fig. S4). We then sought to compare the total fluorescent protein from our images across all conditions. In order to achieve this, cell images were rescaled to our FCS data by rank normalisation as shown by the Q-Q plot (Fig. 4, D) [49].

Since the two datasets shared the same distribution, we calculated the integrated calibrated intensity of GFP fluorescence for each strain, condition and target protein (Fig.4, E and F). We observed that the total Fis1p-GFP detected in *Sc* FIS1^GFP/+^ cells growing in glycerol differed significantly compared to glucose, with a median of 571015 (95% CI: 533732-608517) compared to 467631 (95% CI: 388621-518406) (Fig. 4, E; Man Whitney two-tailed t-test, p<0.0001). Similarly, the median total Fis1p-GFP for *Sc* FIS1^GFP/-^was found to be 1167434 (95% CI: 1118632-1231539) for glycerol and 488846 (95% CI: 471983-512412) for glucose, respectively (Fig. 4, E; Man Whitney two-tailed t-test, p<0.0001). Following the same trend, the arbitrary measurements of total Mdv1p-GFP were significantly different between the two growth conditions for both *Sc* MDV1^GFP/+^ and *Sc* MDV1^GFP/-^ cell populations (Man Whitney two-tailed t-test, p<0.0001; Fig. 4, F). These results suggest that yeast cells growing in non-fermentative media (glycerol) show increased Fis1p and Mdv1p total levels compared to fermentative growth (glucose). As in the case for mitochondrial-associated Mdv1p, no differences were observed in the total cell Mdv1p between the *Sc* MDV1^GFP/+^ and *Sc* MDV1^GFP/-^ strains. In contrast, increased total cell Fis1p was observed in *Sc* FIS1^GFP/-^ cells growing in glycerol but not in glucose compared to *Sc* FIS1^GFP/+^ cells.

### Protein mobility measurement in live *S. cerevisiae* single cells shows an increase of the bound-versus the unbound-state for Mdv1p-GFP mitochondrial associated molecules

To better characterise the dynamics of the mitochondrial-associated Fis1p-GFP and Mdv1p-GFP proteins, we studied their mobility and calculated the proportion between their bound- and unbound-molecules using the set of hemizygote and homozygote *S. cerevisiae* strains created in this study (Table 1). To quantify the kinetics of the proteins, we used a two-component model that allows the estimation of a ‘fast’ diffusion rate of highly mobile molecules and a ‘slow’ diffusion rates of less mobile fractions (see methods). The ‘fast’ diffusion rate represents protein molecules that are free (unbound) in the cytoplasm whereas the ‘slow’ diffusion rate represents protein molecules that are in a bound-state. We found that the majority of Fis1p-GFP molecules diffused fast for measurements taken on mitochondrial locations in *Sc* FIS1^GFP/+^ (n = 127) and *Sc* FIS1^GFP/-^ (n = 80) cells, growing in rich medium (Fig. 5, A-C). Specifically, 59.6% of Fis1p-GFP molecules in *Sc* FIS1^GFP/+^ and 61.2% in *Sc* FIS1^GFP/-^ diffused at a faster pace with an average rate of 12.25±0.69 μm^2^/s (mean±SD) and 9.48±0.65 μm^2^/s (mean±SD), respectively (Fig. 5, A and B). These data suggest that Fis1p-GFP is primarily present on the mitochondria in an unbound protein state. The deletion of one allele is not expected to change the mobility dynamics of the protein. In fact, no differences were observed in the comparison between the slower-moving mobile fractions between *FIS1* homozygote and heterozygote cell populations (Mann-Whitney two-tailed t-test, p > 0.05) (Fig. 5, C).

**Figure 5.**
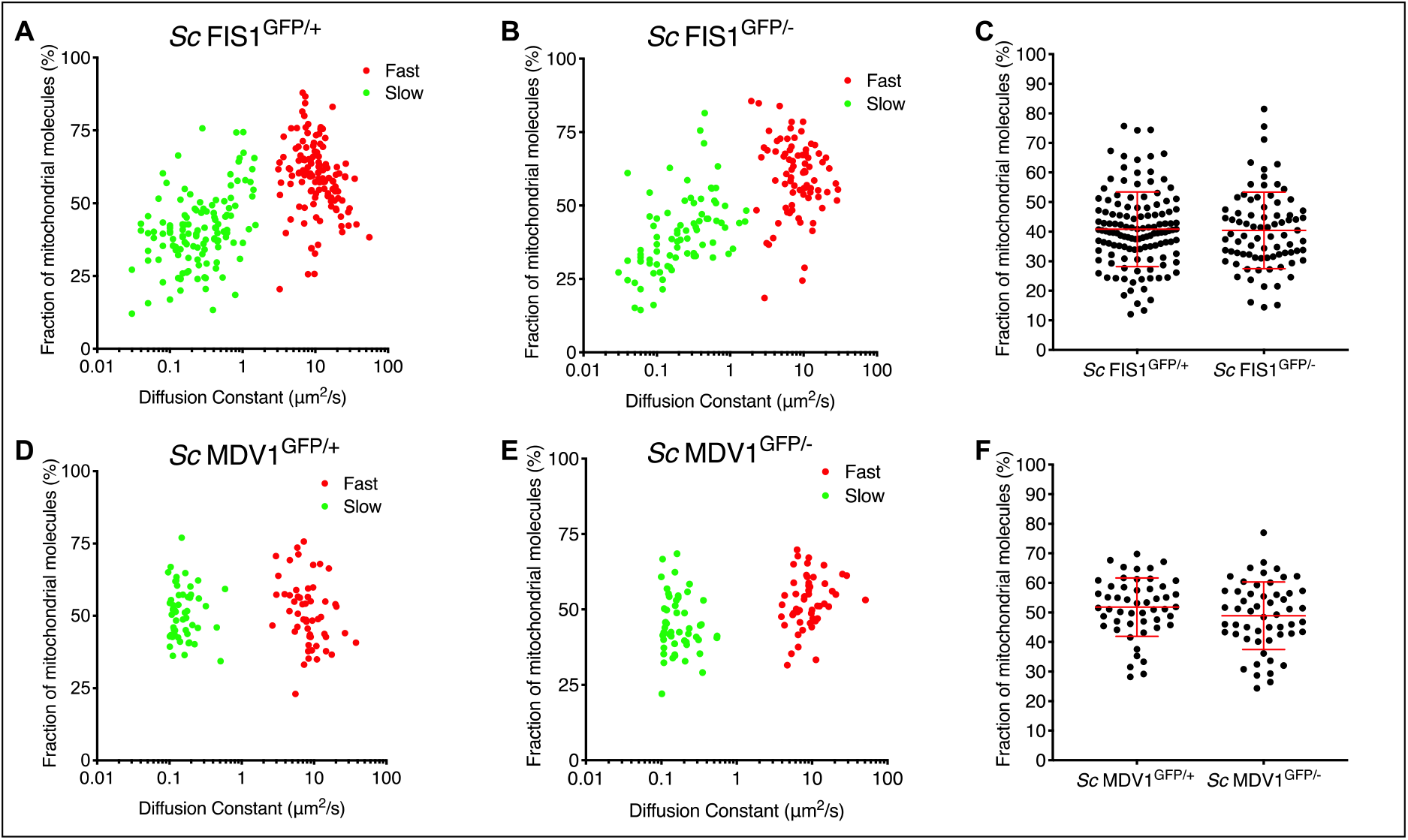
Modelling the mobility kinetics of Fis1p-GFP and Mdv1p-GFP molecules in yeast growing in rich fermentable medium. Mitochondrial Fis1p-GFP was measured in single *Sc* FIS1^GFP/+^ (n = 127) and *Sc* FIS1^GFP/-^ (n = 80) cells growing in media containing glucose. The data obtained on the kinetics of Fis1p-GFP were fitted to a two-component model to estimate the numbers of fast-diffusing (red spots) and slow-diffusing (green spots) molecules. Shown are 2D plots comparing the diffusion rate to the fraction of mobile molecules measured in the confocal volume for the *Sc* FIS1^GFP/+^ (A) and *Sc* FIS1^GFP/-^ (B) cells. The calculated percentage of molecules diffusing slowly was compared between the *Sc* FIS1^GFP/+^ and *Sc* FIS1^GFP/-^ cells using the Mann Whitney two-tailed t-test (C). Mitochondrial Mdv1p-GFP was measured in single *Sc* MDV1^GFP/+^ (n = 54) and *Sc* MDV1^GFP/-^ (n = 51) cells growing in media containing glucose. The data obtained on the kinetics of M1,2dv1p-GFP were fitted to a two-component model to estimate the numbers of fast-diffusing (red dots) and slow-diffusing (green dots) molecules. Shown are 2D plots comparing the diffusion rate to the fraction of mobile molecules measured in the confocal volume for the *Sc* MDV1^GFP/+^ (D) and *Sc* MDV1^GFP/-^ (E) cells (F). The calculated percentage of molecules diffusing slowly was compared between the *Sc* MDV1^GFP/+^ and *Sc* MDV1^GFP/-^ cells using the Mann Whitney two-tailed t-test. Errors bars show mean and SD.

On mitochondrial locations of cells expressing Mdv1p-GFP, the portion of Mdv1p-GFP fast- and slow-moving protein is similar (Fig. 5, D and E). However, still the majority of the mitochondrial-associated Mdv1p-GFP subunits is present in an unbound protein state. We calculated that 51.1% of Mdv1p-GFP in *Sc* MDV1^GFP/+^ (n = 54) and 51.6% in *Sc* MDV1^GFP/-^ (n = 51) molecules diffused at faster pace with an average rate of 9.56±0.83 μm^2^/s (mean±SD) and 10.46±1.08 μm^2^/s (mean±SD), respectively (Fig. 5, D and E). Similarly to Fis1p-GFP, no differences were observed between the slower-moving mobile fractions of the *Sc* MDV1^GFP/+^ and *Sc* MDV1^GFP/-^ cell populations (Mann-Whitney two-tailed t-test, p > 0.05) (Fig. 5, F), indicating that mobility dynamics are independent from the allelic state.

We next investigated whether growth on non-fermentable source, such as glycerol, affects the kinetics of the Fis1p-GFP and Mdv1p-GFP mitochondrial-associated components. We found that the majority of Fis1p moves also fast in YP-glycerol, indicating an unbound state. We calculated the mobility dynamics of Fis1p-GFP to be at an average rate of 11.11±0.73 μm^2^/s (mean±SD) for 58.5% of molecules in *Sc* FIS1^GFP/+^ cells (n = 80) and of 11.09±0.86 μm^2^/s (mean±SD) for 62% of molecules in *Sc* FIS1^GFP/-^ cells (n = 70) (Fig. 6, A and B). On the contrary, the proportion of the slow-moving mitochondrial-associated Mdv1p-GFP fractions is higher compared to the fast-moving molecules in *Sc* MDV1^GFP/+^ (n = 51) and *Sc* MDV1^GFP/-^ (n = 48) cells (Fig. 6, D and E). We calculated that 55.5% of *Sc* MDV1^GFP/+^ and 58.4% of *Sc* MDV1^GFP/-^ molecules diffused at a slower rate with an average rate of 0.15±0.01 μm^2^/s (mean±SD) and 0.18±0.01 μm^2^/s (mean±SD), respectively. We observed no significant differences between the slower-moving mobile fractions of the *Sc* FIS1^GFP/+^ and *Sc* FIS1^GFP/-^ cell populations and the *Sc* MDV1^GFP/+^ and *Sc* MDV1^GFP/-^ (Mann Whitney two-tailed t-test, p>0.05) (Fig. 6, C and F), indicating that the fission proteins show similar mobility dynamics in both cell types. Overall, the above data suggest that the bound-state of the mitochondrial Fis1p in *Sc* FIS1^GFP/+^ and *Sc* FIS1^GFP/-^ cells is not affected by the deletion of one *FIS1* allele, either in glucose or glycerol media. On the other hand Mdv1p bound-state appears to increase upon growth on non-fermentable carbon sources.

**Figure 6.**
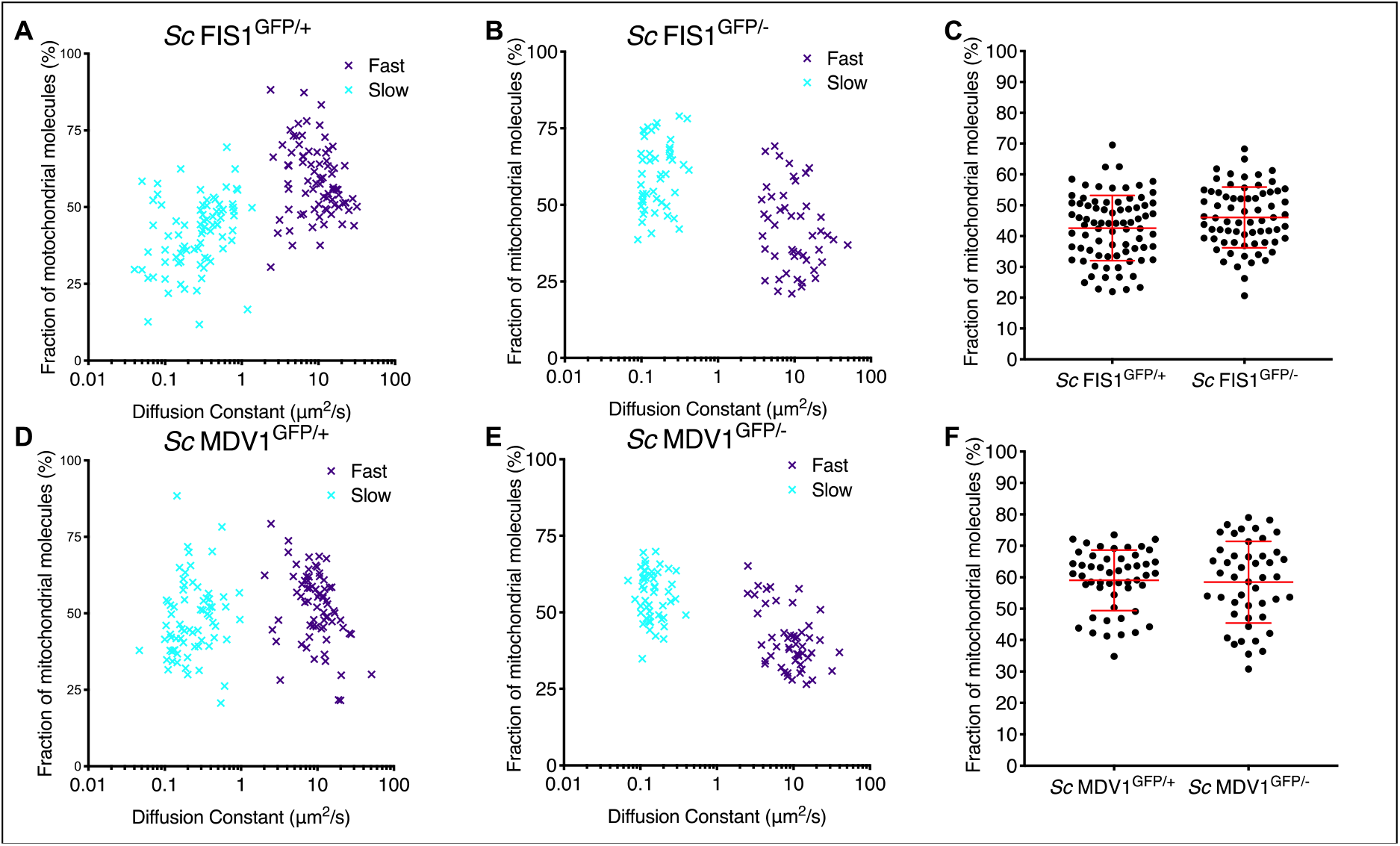
Modelling Fis1p mobility in respiring GFP-labelled living yeast cells by FCS. The FCS data on mitochondrial Fis1p-GFP obtained for single *Sc* FIS1^GFP/+^ (n = 80), *Sc* FIS1^GFP/-^ (n = 70), *Sc* MDV1^GFP/+^ (n = 51), *Sc* MDV1^GFP/-^ (n = 48) cells growing in media containing glycerol were fitted to a two-component model to estimate the numbers of fast-diffusing (purple spots) and slow-diffusing (light blue spots) molecules. Shown are 2D plots comparing the diffusion rate to the fraction of mobile molecules measured in the confocal volume for the *Sc* FIS1^GFP/+^ (A), *Sc* FIS1^GFP/-^ (B), *Sc* MDV1^GFP/+^ (D) and *Sc* MDV1^GFP/-^ (E) cells. Scatter plots of the proportion of slow-moving Fis1p-GFP molecules in *Sc* FIS1^GFP/+^ and *Sc* FIS1^GFP/-^ cells are shown in panel (C) and *Sc* MDV1^GFP/+^ and *Sc* MDV1^GFP/-^ cells in panel (F). Statistical analysis was performed using the Mann-Whitney two-tailed t-test. Errors bars show mean and SD.

By comparing the diffusion constants of the Fis1p-GFP and Mdv1-GFP in cells growing in glucose versus glycerol, we were able to analyse how the environmental condition affects the mobility dynamics of each protein (Fig. 7). While in *Sc* FIS1^GFP/+^ cells the fraction of slow-moving protein molecules remains the same in the two media conditions (Fig. 7, A), in the *Sc* FIS1^GFP/-^ background, it increases significantly under strict respiration (Mann Whitney two-tailed t-test; p = 0.0001) (Fig. 7, B). Functional mitochondria are essential for the efficiency of yeast cells growing in glycerol. Therefore, this result could indicate that in the case where only one *FIS1* allele is expressed in the cell, a higher number of the endogenous Fis1p binds to mitochondria in order to ensure the existence of active mitochondrial structures and ultimately, the maintenance of the mitochondrial function under respiration. Compared to the glucose medium (Fig. 7, C and D), the fraction of slow-moving Mdv1p was significantly increased in both the *Sc* MDV1^GFP/+^ (Mann Whitney two-tailed t-test; p = 0.0151) and *Sc* MDV1^GFP/-^ (Mann Whitney two-tailed t-test; p = 0.0030) under strict respiration conditions (Fig. 7, C and D). Collectively, these data suggest that the abundance (see Fig. 4, B) and the bound-state of the Mdv1p (Fig. 7, C and D) are drastically affected by the media conditions.

**Figure 7.**
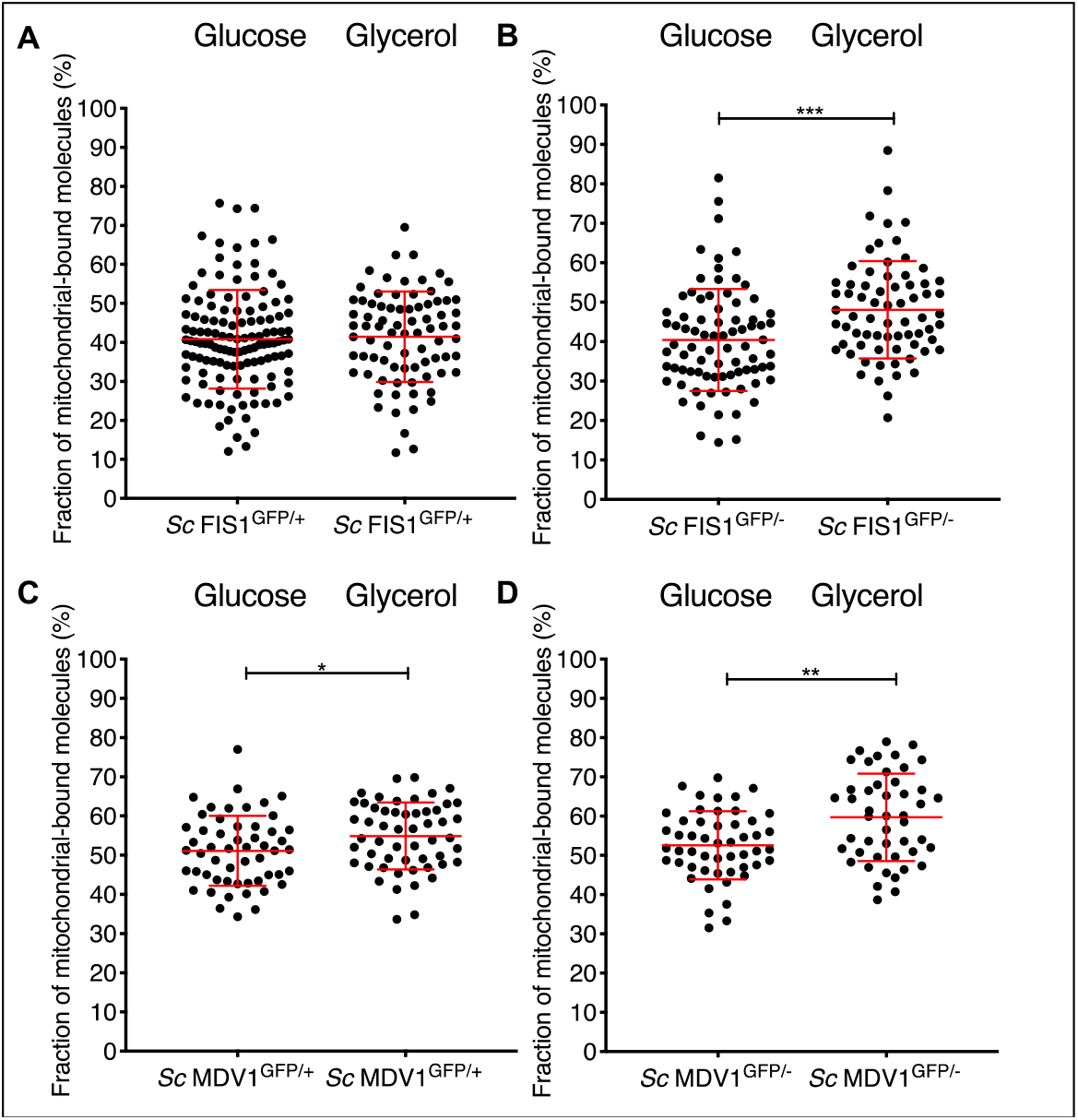
Comparison of the diffusion constants of Fis1p-GFP and Mdv1p-GFP between *S. cerevisiae* cells growing in fermentable and non-fermentable carbon sources. Scatter plots of Fis1p-GFP diffusion rates for the fractions of slowly moving molecules in individual *Sc* FIS1^GFP/+^ (A) and *Sc* FIS1^GFP/-^ (B) cells growing in glucose (*Sc* FIS1^GFP/+^ n = 127, *Sc* FIS1^GFP/-^ n = 80) or in glycerol (*Sc* FIS1^GFP/+^ n = 80, *Sc* FIS1^GFP/-^ n = 70). Scatter plots of Mdv1p-GFP diffusion rates for the fractions of slowly moving molecules in individual *Sc* MDV1^GFP/+^ (C) and *Sc* MDV1^GFP/-^ (D) cells growing in glucose (*Sc* MDV1^GFP/+^ n = 54, *Sc* MDV1^GFP/-^ n = 51) or in glycerol (*Sc* MDV1^GFP/+^ n = 54, *Sc* MDV1^GFP/-^ n = 47). *** represents significance for p ≤ 0.001 for Mann Whitney two-tailed t-test.

## Discussion

In yeast, the nuclear-encoded Fis1p and Mdv1p have been identified as essential components of the mitochondrial fission machinery while their recruitment to the mitochondrial membrane is of vital importance not only for the scission of mitochondria, but also for the retention of functional forms of the organelles [31, 50, 51]. At present, 13 independent large-scale studies on the *S. cerevisiae* yeast provide different results on protein concentrations for Fis1p and Mdv1p and six of those provide additional information on the localisation of the proteins [6, 7, 9, 25–29, 52–56]. Here we optimised the fluorescence spectroscopy techniques, FCS, to precisely quantify the native mitochondrial and cellular expression of the low abundant Fis1p and Mdv1p fission proteins and compare the absolute protein copy numbers and mobility states over allelic expressional modifications, and in different environmental conditions.

We found that GFP-fused Fis1p was significantly higher on mitochondrial locations of the *Sc* FIS1^GFP/-^ cells compared to *Sc* FIS1^GFP/+^, indicating higher protein abundance in the strains lacking one *FIS1* allele compared to the strains having both alleles under both fermentative and respiratory growth (Fig. 2, D; Fig. 3, D). Since *FIS1* gene is haplo-insufficient in carbon-limited medium [57, 58], it is possible that the cell may try to compensate the protein levels of the remaining *FIS1* allele to counteract the reduced growth rates [57, 58]. Beside mitochondrial fission, Fis1p also participates in multiple mitochondrial-related activities, including tethering of damaged or misfolded proteins to mother cells [31], regulation of ethanol-induced apoptosis [32, 33] and tuning of cell death in aging cells [34]. Thus, the maintenance of mitochondrial-associated Fis1p at sufficient levels is key for the cellular growth and integrity. Importantly, cells lacking one *FIS1* allele showed similar compensation in the total cellular Fis1p when grown in glycerol, but not in glucose, compared to those having both alleles (Fig. 4, E). Notably, the highest amount of total Fis1p was observed in *Sc* FIS1^GFP/-^ cells grown in glycerol, with a two-fold difference from all other conditions (Fig. 4, E), supporting the suggestion that Fis1p plays an important role in respiring mitochondria. It is known that functional mitochondria are essential for respiratory growth in yeast, and thus, under these conditions cells may need to compensate the lack of one copy of *FIS1* by producing more protein compared to the glucose-rich condition where the metabolic requirements are met mainly by fermentation.

Mdv1p did not show a compensatory trend when quantified in *Sc* MDV1^GFP/+^ and *Sc* MDV1^GFP/-^ strains growing either in glucose (Fig. 2, I) or in glycerol (Fig. 3, I). These data suggest that the protein levels of Mdv1p associated to mitochondria are not influenced by the deletion of the *MDV1* allele. The total cellular Mdv1p was neither increased nor decreased in strains carrying only one *MDV1* allele under any growth condition (Fig. 4, F). In contrast to *FIS1* allele, *MDV1* is not associated to neither a haplo-insufficient or haplo-proficient phenotype when in hemizygous state [57, 58]. Moreover, the localisation of Mdv1p to mitochondria is exclusively related to the regulation of the fission process, and our data indicate that the amount of Mdv1p protein from one allelic locus is probably able to provide the stoichiometric balance required for the correct formation of the Mdv1p/Caf4p::Fis1p::Dnm1p protein complex in diploid *S. cerevisiae* strains.

In both cases of Mdv1p and Fis1p it appears that the quantitative perturbation of the protein abundance is not reflected to the bound state of the proteins in cells growing under fermentation or respiration. This finding comes in support of the proposed hypothesis that the two proteins act in conjunction in order to fulfil their functional roles in the mitochondrion [48]. Accordingly, it could be suggested that an alteration in the quantity of one of the two interacting proteins might need an alteration in the quantity of the other in order to maintain the stoichiometry of the complexes that they are part of.

We observed that growth under respiratory conditions exerted a strong effect on the cellular Fis1p and Mdv1p abundance. Both homozygote and heterozygote strains demonstrated significant increase in cellular Fis1p and Mdv1p when grown in glycerol compared to growth in glucose (Fig. 4, E and F). This is in accordance with previous reports showing an increase of abundance of mitochondrial proteins when cells are shifted from fermentable (glucose) to the non-fermentable carbon source (glycerol) [29].

Moreover, environmental conditions can influence the size of the mitochondrial network, hence the abundance of its associated proteins. Specifically, it has been shown that when *S. cerevisiae* is grown on medium containing glycerol as the only carbon source the cells exhibit a strongly branched tubular reticulum compared to glucose-grown cells [59]. In our study, we quantified mitochondrial surface in positive cells showing mitochondrial GFP fluorescence in comparison to negative control cells carrying Cox4p-DsRed-stained mitochondria (Fig. 4, C; Fig. S3), and confirmed that the mitochondrial mass increases in cell grown in glycerol (Fig. S3) in agreement with previous studies [59]. Here, we showed that under respiration the mitochondrial associated Mdv1p increase, while the Fis1p does not (Fig. 4). This result indicates that the growth conditions have a differential effect on the abundance of Mdv1p and Fis1p associated to mitochondria. This could be explained because while the only known biological function of Mdv1p is mitochondrial and peroxisome fission, hence requires mitochondria association of Mdv1p, Fis1p has a more ambiguous role, since the protein is found in different cell compartments [29] and is involved in other functions such as apoptosis [29].

The majority of the mitochondrial-associated Fis1p-GFP copies in *Sc* FIS1^GFP/+^ cells were found to be at an unbound protein state under both fermentative and respiratory conditions, with no significant differences in the portion of free copies between the two environments (Fig. 5, A; Fig. 6, A). However, the fraction of bound molecules was significantly increased in the *Sc* FIS1^GFP/-^ cells growing under respiration (~48%) compared to the *Sc* FIS1^GFP/-^ cells growing under fermentation (~40%), even though the abundance of the Fis1p remained the same (Fig. 4, A). This result could indicate that in respiring yeasts a higher portion of the Fis1p copies are in bound state in order to ensure the existence of functional mitochondria and ultimately, the efficiency of the cells [60]. Significant changes were observed in the bound state of mitochondrial Mdv1p molecules of both *Sc* MDV1^GFP/+^ and *Sc* MDV1^GFP/-^ cells growing under respiration compared to fermentation. This is in concordance with the significant increase in the mitochondrial Mdv1p abundance in *Sc* MDV1^GFP/+^ and *Sc* MDV1^GFP/-^ cells growing under respiratory conditions. This could be anticipated based on the known requirement for increased fission when cells are grown under respiratory conditions [60] and the role of Mdv1p in this process, where more Mdv1p molecules are to participate in a heightened number of Dnm1p-containing protein complexes that regulate fission [31, 61].

This study shows the power of using FCS in combination with imaging analysis to quantify low abundant proteins, which are difficult to detect with other analytical techniques [8, 9]. Besides quantifying the variation in the abundance of low-expressed proteins, their location and mobility rates can be also assessed. All together these factors can ultimately affect the phenotypic response of the yeast cell to different environmental stimuli. Collectively, our data support a differential ability of Fis1p and Mdv1p to buffer copy number variation and shows the different effects that the environment have on the abundance of these proteins in the mitochondria, including their altered mobility states. The ability to generate quantitative parameters, such as kinetic rates and protein concentration allows to gain a deeper understanding into the dynamics that dictate the biological role of low-abundant proteins. Furthermore, being able to detect fluctuations in protein levels after allelic variation in different environmental contexts will help to determine the level of robustness and the degree of plasticity of these proteins.

## Experimental procedures

### Yeast strains and plasmids

The *S. cerevisiae* BY4741 (ATCC^®^ 201388TM), BY4742 (ATCC^®^ 201389TM) and BY4743 (ATTCC^®^ 201390TM) strains were obtained from the American Type Culture Collection. The *S. cerevisiae* BY4741 FIS1^GFP^ (Clone ID: YIL065C) and *S. cerevisiae* BY4741 MDV1^GFP^ (Clone ID: YJL112W) fusion strains were purchased from Thermo Fisher Scientific (Cat. No. 95700). Yeast strains were routinely maintained on solid Yeast Peptone Dextrose (YPD) (2% (w/v) BactoTM Peptone; 1% (w/v) BactoTM yeast extract; 2% (w/v) glucose) media containing 2% (w/v) agar and plates were stored at 4°C.

### Yeast transformation and mating type switching

Yeasts were transformed via chemical methods as previously described [66]. The homothallic *S. cerevisiae* BY4741 *MAT*a GFP-labelled strains were subjected to mating type switching as previously described in Amberg *et al*. [67]. Haploid *S. cerevisiae* strains subjected to mating type switch were checked for their mating type by colony PCR for the *MAT* locus (Table S1).

### Diploid strain construction by single-cell mating

Single cells of fluorescently-labelled *S. cerevisiae MAT*a strains were manually crossed with *MAT*α cells of the same strain background (a) carrying no fluorescent markers, (b) having the *FIS1* or *MDV1* genes deleted or (c) containing the same fluorophore tags, using the Singer Instruments (MSM-300) micromanipulator. Colonies of *S. cerevisiae* strains engineered either to contain one allelic copy fused to a fluorophore and one wild-type copy (*Sc* FIS1^GFP/+^ or *Sc* MDV1^GFP/+^) or only the fluorescently-fused allele (*Sc* FIS1^GFP/-^ or *Sc* MDV1^GFP/-^), were selected based on auxotrophic markers and replica-plated on minimal SD medium supplemented with 0.2% (w/v) uracil, 0.1% (w/v) leucine and 2% (w/v) agar. The double-tagged diploids, which contained two different antibiotic markers, were selected on YPD agar plates supplemented with the antibiotics G418 and hygromycin B. The ploidy of the generated diploids was checked by *MAT* locus colony PCR (Table S1).

### Ploidy analysis by Fluorescence-Activated Cell Sorting (FACS)

FACS analysis was carried out as previously described [68]. The stained cells were analysed at 488 nm excitation and 523 nm emission using a BD LSR Fortessa cell analyser (BD Biosciences, USA) from the Flow cytometry core facilities of the University of Manchester (UK).

### Strain preparation for live cell imaging and FCS

Cells grown under fermentative conditions: diploid *MAT**α***/α strains for imaging were inoculated into 5 ml of YPD medium and grown overnight. Saturated cultures were diluted into 5 ml of fresh medium and grown to an D_600_ of 0.5. The optical density of cultures was calculated using a Jenway Genova spectrophotometer (Bibby Scientific, UK). Cells were collected by centrifugation at 4000 rpm for 3 minutes, washed twice with sterile MilliQ water and diluted into 500 μl of fresh YPD in 35 mm glass-bottomed CELLview^™^ microscopy dishes (Greiner Bio-One, UK). Following a step-by-step optimization procedure, cells at the mid log phase were imaged in minimal SD media supplemented with essential amino acids. During this process, cells were seeded into individual wells coated with the cell adhesive Corning^®^ Cell-Tak (Sigma, Prod. no. # DLW354241) in a final concentration of 5 μg/μm^2^. Prior to cell transfer, 250 μl of cell adhesive solution (25 μl Corning Cell-Tak and 725 μl of Sodium bicarbonate pH 8.0) was added to each compartment, incubated at room temperature (RT) for two hours and washed once with sterile MilliQ water.

Cells grown under respiratory conditions: *S. cerevisiae* strains expressing the GFP-fusion proteins were inoculated into 5 ml of YP medium containing 2% glycerol (YPgly) and grown overnight. Saturated cultures were diluted into 5 ml of fresh medium and grown to an D_600_ of 0.5. The optical density of cultures was calculated using a Jenway Genova spectrophotometer (Bibby Scientific, UK). Cells were collected by centrifugation at 4000 rpm for 3 minutes, washed twice with sterile MilliQ water and diluted into 500 μl of minimal SD medium containing 2% glycerol and supplemented with the required amino acids. Cells were seeded into coated 35 mm glass-bottomed Greiner dishes and imaged at 30°C with no additional CO2. Strains on the same plate were imaged consecutively with 30-minute intervals in a random sequence. All confocal microscopy results illustrated in this work represent imaging and FCS data collected from single live cells over a set of four to six independent experiments per condition and yeast strain.

### Confocal microscopy and live cell imaging

A Zeiss LSM880 microscopy system with GaAsP detectors attached to the inverted Axio observer Z1 microscope with a 63x C-apochromat, 1.4 NA oil immersion objective was used. Excitation of GFP was performed using an Argon ion laser at 488 nm. Emitted light was detected between 490 and 552 nm. DsRed was excited at 561 nm and emission detected between 567 and 614 nm. These emission wavelengths were selected as the optimum to avoid any spillover between fluorophores and to avoid autofluorescence as assessed by lambda scans.

### Lambda scanning

The lambda scan function of the Zen 2010B Zeiss software using both 488 and 561 nm excitation gave 10 simultaneous images in 10 nm steps from 498-687 nm which were combined in a single image. These spectral images were then analysed by linear unmixing to generate distinct emission profiles for each probe across the wavelength range and discriminate the contribution of the individual fluorescent proteins and the contribution of autofluorescence.

### Protein quantification by FCS

FCS was carried out using the same excitation and emission strategy with reduced laser power to minimise photobleaching. Fluorescence fluctuation counts of a minimum 0.5 counts per molecule were collected through a pinhole set to one Airy unit. Photon counts were recorded for 10 seconds and 10 repetitions for each measurement or adjusted to 5X5 second runs for the more sensitive detectors as outlined in Kim *et al*. [12]. Mean fluorescence intensities of GFP fusion proteins were calculated from measurements obtained either from cytoplasmic or mitochondrial fluorescence emissions for free and bound molecules, with a binning time of 200 ns. The FCS measurements obtained from the protein quantification experiments were calculated automatically into their autocorrelation functions using the Zeiss-built-in ZEN 2010B software. Protein mobility measurements were recorded and analysed manually using the freely available PyCorrFit software [49].

### Statistical Analysis of protein quantification data

Analysis of fluorescence correlation data was carried out in Microsoft Excel and GraphPad Prism, version 7.0e for Mac OS C (www.graphpad.com). As a first step of analysis, parameters were fitted manually to a two-component free diffusion model using the Zen 2010B Zeiss software. This allowed the removal of measurements that gave abnormally small or large values for the non-fixed structural parameter, which were defined as below 0.1 and above 15, respectively. The remaining measurements were then fitted to a two-component diffusion model using a fixed structural parameter at 4. The unfit measurements with chi^2^ values less than 10^-4^ were considered to describe unsuccessful fits between the model function and the experimental data and therefore, were excluded from further analysis. Finally, in the generated correlation curves, the first five signal recordings of the confocal area were also removed as these frames typically encompass a ~0.5-second period of non-correlated background signals, such as noise.

The analysis of fitted data to obtain the number of amplitude molecules and the counts per second per molecule was performed manually in Microsoft Excel. For each measurement, the data collected per single cell throughout the repetition runs (10X10 or 5X5 runs) were averaged and copied into a GraphPad Prism file to generate the appropriate graph plots and calculate the standard deviation (SD) values and 95% confidence intervals of the selected data sets (column statistics function). Values were tested for normal distribution using D’Agostino-Pearson omnibus K2 normality test. Statistical comparison between two groups was performed using the non-parametric Mann Whitney two-tailed t-test for non-normally distributed measurements. Linear regression was used to monitor the relation between the CPM and amplitude measurements. Goodness of fit was evaluated using the R^2^ statistic measure.

### Measuring protein mobility

To calculate the diffusion rates *D_i_* of the fluorescent molecules, the experimental correlation data obtained per confocal volume by FCS were fitted into a two-component 3D diffusion model with a triple state component [49, 62–64], using the PyCorrFit software [49].

### Statistical Analysis of diffusion rates

The analysis of fitted data to obtain the diffusion constants *D_1_* and *D_2_* of the fluorescent particles detected in the confocal volume was performed manually in Microsoft Excel. The lateral beam of the confocal volume (*w_0_*) dimensions was previously estimated to be 0.22±0.063 μm using Rhodamine 6G (Systems Microscopy Center, University of Manchester). The diffusion constant *D_1_* was calculated in relevance to the characteristic diffusion time *τ_1_* (μs) measured for the *n_1_* fraction of the detected molecules (*D_1_* = *w_0_^2^* / 4*τ_1_*). Similarly, the diffusion constant *D_2_* was calculated in relevance to the characteristic diffusion time *τ_2_* (μs) measured for the *n*_2_ fraction of the detected molecules (*D_2_* = *w_0_^2^* / 4*τ_2_*).

The quality of the generated data was checked following a similar pipeline to the protein quantification data analysis protocol. Unfit measurements with chi^2^ values less than 10^-4^ were considered as indicative of poor model fitting and thus, were excluded from further analysis. The noise spectrum was removed by starting the analysis of the fitted curve from timepoint 0.5 (seconds) to either timepoint 10 (seconds) or 5 (seconds) based on the duration of the signal measurements. For each measurement, the diffusion constant data collected per single cell throughout the repetition runs (10X10 or 5X5 runs) were averaged for a minimum of 5 or 3 repetitions, respectively.

Finally, the parameter values of the diffusing molecular fractions *n_1_* and *n_2_*, calculated in proportion to the total number of the diffusion molecules *n* detected in the confocal volume, and the respective diffusion constants *D_1_* and *D_2_*, were fit in a linear-regression model using the GraphPad Prism program (version 7.0e for Mac OS C, www.graphpad.com). The percentage of variance among the fitted values obtained from experiments carried out under the same conditions was calculated by the R^2^ goodness-of-fit statistic measure. Values were tested for normal distribution using D’Agostino-Pearson omnibus K2 normality test. Statistical comparison between two groups of normally distributed mean measurements was performed using the parametric unpaired t-test.

### Quantification of cell size and mitochondrial surface

‘Segmentation and quantification of subcellular shapes’ (SQUASSH) image J plugin was used to segment and identify mitochondria from images of individual cells that were automatically cropped from whole field images [49]. In order to do this, individual cells from confocal microscopy images were identified using Cell Profiler. In brief, the image was subjected to a blur processing stage to enhance large objects. Then objects with a range of pixels from 10 to 200 were subsequently identified using the ‘adaptive thresholding’ function. Examples of individual cropped cells shown in Supplementary Figure 4 (Fig. S4). For each condition, several hundred to a few thousand cells were isolated and a subset fraction of 100 cells with a size between 10 - 15 μm^2^ were randomly chosen using a custom MATLAB script. The selected data sets were then analysed by SQUASSH (parameters: rolling ball window = 10, regularization = 0.05, minimum intensity = 0.15, noise model = poisson and objects below 2 pixels removed). The SQUASSH plugin exports various data outputs pertaining to the segmented space, including mitochondrial footprint (the area of the segmentation) and the branch characteristics (length and branching of rod-like structures). These were converted from values in pixels to μm^2^ using the recorded pixel scale from the original confocal images.

### Total RNA extraction and quantitative RT-PCR

Triplicate cultures (25 mL) of each strain were grown in YPD to an OD_600_ of 0.5 (mid-log phase). The cells were then harvested by centrifugation at 3000 g for 1 minute, washed in 2 mL of sterile H2O, and frozen with liquid nitrogen. Total RNA was isolated using RNeasy Mini Kit (Qiagen) following the protocol for enzymatic digestion of cell wall followed by lysis of spheroplasts. The quality and concentration of RNA were determined by a NanoDrop (Thermo Scientific). An A260/A230 ratio > 2 and an A260/A280 ratio in the range of 1.8–2.2 were considered acceptable. One microgram of total RNA was reverse transcribed into cDNA using QuantiTect Reverse Transcription Kit (Qiagen) following the manufacturer’s protocol. QuantiTect Reverse Transcription Kit includes a genomic DNA removal step. However, the absence of contaminant genomic DNA in RNA preparations was verified using RNA as a template in the real-time PCR assay. The signals obtained represent the expression levels of the GFP-tagged *FIS1* and *MDV1* and were normalized to that of Actin (ACT1) and quantified by the ΔΔCt method [65]. For each condition, the resulting expression levels are presented as the mean ± SD of three independent experiments, each performed in triplicate and all runs included a no template control (NTC), and a control lacking reverse transcriptase (-RT) (Fig. S2).

## Data availability

All data described in the manuscript are contained within the manuscript. Cell profiler pipeline, custom MATLAB scripts for mitochondrial and protein mass analysis, and fluorescently tagged strains are available on request.

## Acknowledgments

We are grateful to Dr. Spyridon Megremis for his comments, suggestions and proofreading of the manuscript.

## Funding and additional information

AM is supported by a studentship from Biotechnology and Biological Sciences Research Council (BBSRC; BB/M011208/1) under the Doctoral Training Partnership scheme (DTP). LNB is supported by BBSRC grant (BB/T002123/1) to DD.

