## Supplementary Information for "Quantitative characterisation of low abundant yeast mitochondrial proteins reveals compensation for haplo-insufficiency in different environments"

Transformed cells for GFP were subjected to confirmation PCR to verify the integration of the C-terminal tags at the 3' end of *FIS1* or *MDV1* ORFs. Confirmation primers were designed to amplify the upstream and downstream regions of the coding gene and the recombination site specific to the GFP-*His3MX6* cassette. For each strain, the reverse sequence for the upstream confirmation of the tag insert is specific to the cassette and is identical in both strain backgrounds. Similarly, the forward primer for the downstream confirmation of the histidine marker integration is kept the same for all strains. The GFP fusion was confirmed using the genomic DNA of the transformed colonies and the high fidelity LongAmp *Taq* DNA polymerase (New England Biolabs, UK) in a PCR reaction of 25µl. The confirmation primers designed for the *S. cerevisiae* fusion strains are listed in Table S1.

Haploid *S. cerevisiae* mating type was confirmed by colony PCR for the *MAT* locus. Confirmation primers of universal sequences were designed to amplify the *MAT*a and *MAT*α region in *S. cerevisiae* strains (Supplementary Table 1). Colonies of each strain background were suspended in 50µl sterile MilliQ water and heated at 95°C for 15 minutes, and 5µl of the cell suspension were used as template in a 25µl PCR reaction using the MyTaq Red DNA mix (Bioline, UK). PCR conditions were: 94°C for 1 minute, 54°C for 1 minute, 72°C for 1 minute and 72°C for 10 minutes for 30 cycles. To confirm that the constructed GFP fusion strains contained a diploid genome, the DNA content of the examined strains was also analysed by FACS and compared to the *S. cerevisiae* BY4741 haploid and the *S. cerevisiae* BY4743 diploid control strains of known ploidy (Fig. S1).

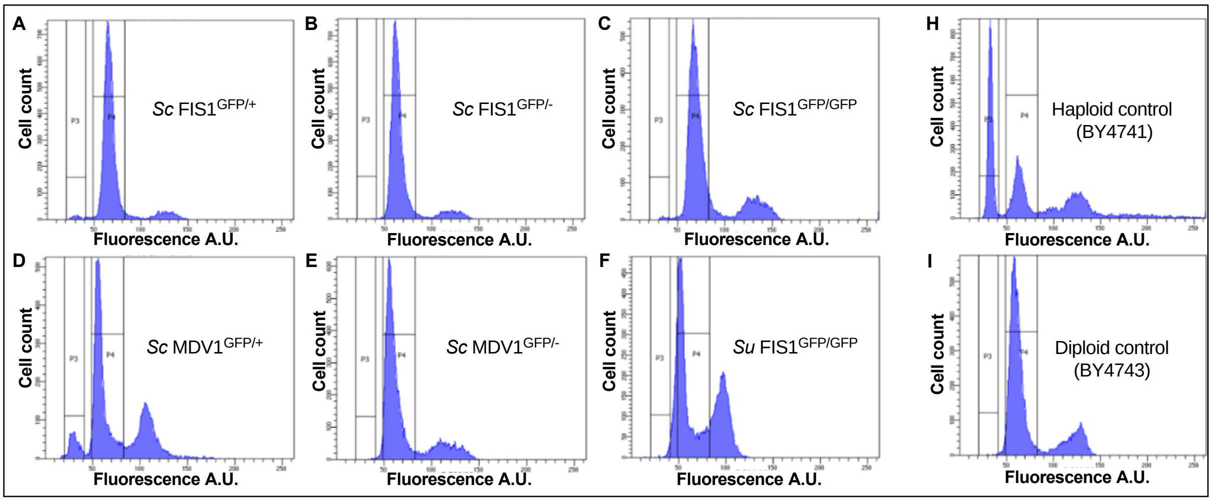


**Supplementary Figure 1. Analysis of ploidy in the GFP *S. cerevisiae* fusion strains.** Flow cytometry analysis of the DNA content of the *FIS1*$^{GFP}$ (A), (B), (C) and *MDV1*$^{GFP}$ (D), (E), (F) *S. cerevisiae* diploid strains in comparison to the BY4741 (haploid control) (H) and BY4743 (diploid control) standards (I). Histograms represent SYTOX® Green fluorescence intensity (A.U.) versus cell count. BY4741 ( Mean A.U. 73.7), BY4743 (Mean A.U. 82.6), *Sc* FIS1$^{GFP/+}$ (Mean A.U. 92.3), *Sc* FIS1$^{GFP/-}$ (Mean A.U. 91.8), *Sc* FIS1$^{GFP/GFP}$ (Mean A.U. 84), *Sc* MDV1$^{GFP/+}$ (Mean A.U. 86.8), *Sc* MDV1$^{GFP/-}$ (Mean A.U. 78.8), *Sc* MDV1$^{GFP/GFP}$ (Mean A.U. 97.4); A.U. represents Arbitrary Units. *Sc* represents *Saccharomyces cerevisiae*.

Primers for quantitative RT-PCR were designed to produce an amplicon between 80–150 bp (Supplementary Table 1). Optimised qPCR reactions contained 5 ng/µl of cDNA, 4 pmol each primer and 5 µL of iTAq Universal SYBR Green super Mix 2X (Bio-Rad) in a final volume of 10 µL. The amplifications were performed on a Light Cycler 480 real time System (Roche) for 35 cycles of: 15 seconds at 95°C; 30 seconds at 57°C; and 30 seconds at 72°C. Melting curve data were collected incrementing by 0.5˚C the temperature from 65˚C to 95˚C using default settings in the machine.

| **Primer name** | **Sequence (5' - 3')** | **Tm°** | **Use in the study** |
|---|---|---|---|
| MAT**a**_locus_F1 | ACTCCACTTCAAGTAAGAGTTTG | 58.1 | Amplification of *MAT* locus for mating type switch |
| MATalpha_locus_F2 | GCACGGAATATGGGACTACTTCG | 67.3 | |
| MAT_locus_R | AGTCACATCAAGATCGTTTATGG | 61.7 | |
| FIS1_Conf_up_F | CCCGTAGACGAGAATGCCTA | 64.0 | Upstream confirmation of GFP-*His3MX6* cassette insertion at the 3' end of *FIS1* |
| FIS1_Conf_up_GFP_R | TGACTTCAGCACGTGTCTTGT | 63.6 | |
| FIS1_Conf_dw_GFP_F | GACGGCCCTATGCTGTTATC | 63.3 | Downstream confirmation of GFP-His3MX6 cassette insertion at the 3' end of *FIS1* |
| FIS1_Conf_dw_R | CCATCTTGTCGTTTGGTCCT | 63.9 | |
| MDV1_Conf_up_F | TGCCTGGTCACAGGTTCATA | 63.4 | Upstream confirmation of GFP-His3MX6 cassette insertion at the 3' end of *MDV1* |
| MDV1_Conf_up_GFP_R | TGACTTCAGCACGTGTCTYGY | 63.6 | |
| MDV1_Conf_dw_F | GACGGCCCTATGCTGTTATC | 63.3 | Downstream confirmation of GFP-His3MX6 cassette insertion at the 3' end of *MDV1* |
| MDV1_Conf_dw_GFP_R | TTTCGACAATTTGGCATCTG | 63.8 | |
| FIS1_GFP_F | AGTCAAGGTTTAACTACGCATGG | 57.0 | Amplification of *FIS1*-GFP for Real-Time PCR |
| FIS1_GFP_R | ACTCGGCCTCTTTGTAAATGTCT | 59.0 | |

**Supplementary Table 1.** Primer sequences for MAT locus amplification; confirmation of GFP fusion and Real-time PCR . Tm°: Melting temperature, F: Forward, R: Reverse.

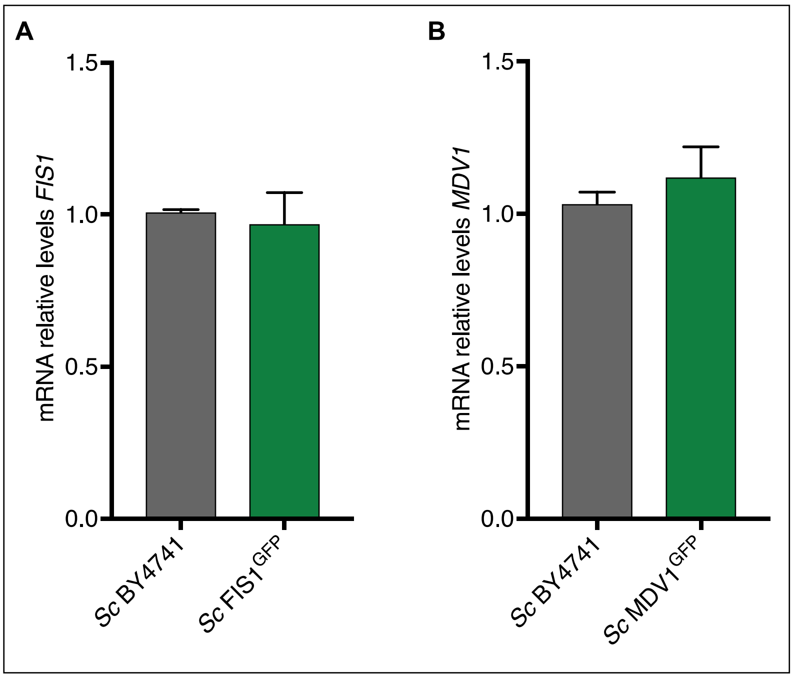


**Supplementary Figure 2. mRNA levels of *FIS1* and *MDV1* are not altered in GFP-tagged strains.** Relative mRNA levels of (A) *FIS1* and (B) *MDV1* analysed by RT-qPCR in the wild-type and in the GFP-tagged *Sc* $FIS1^{GFP}$ and *Sc* $MDV1^{GFP}$ haploid strains, respectively; error bars represent standard deviation of three biological replicates; *Sc* indicates *Saccharomyces cerevisiae*. P-values from significance testing were calculated using unpaired Student's T-test.

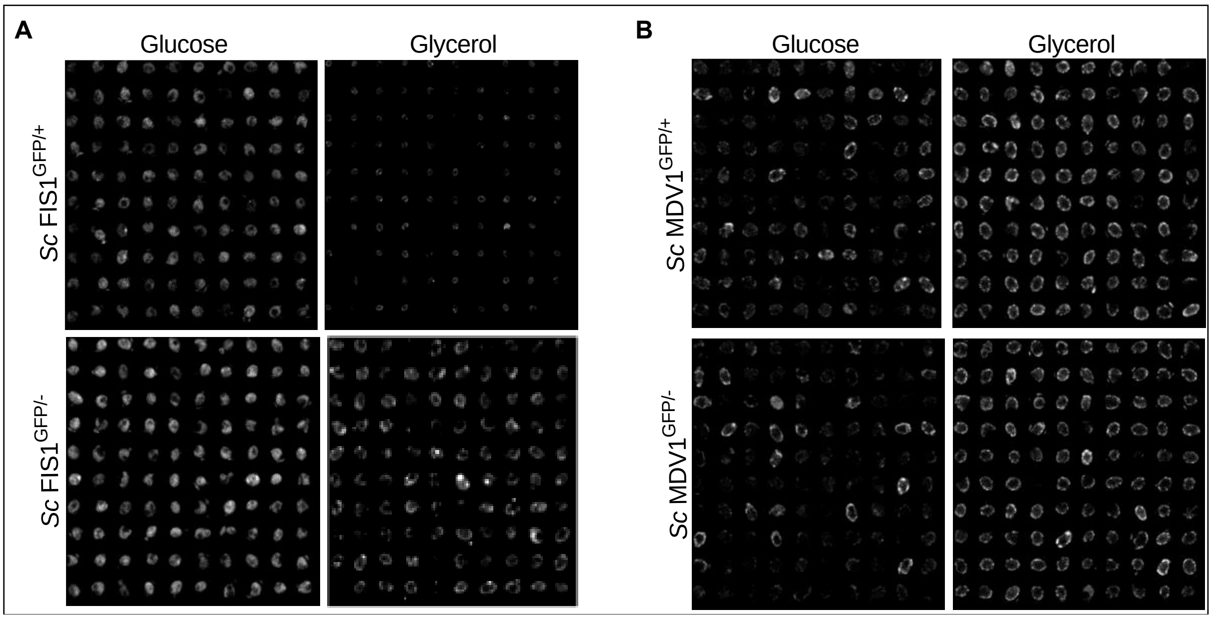


**Supplementary Figure 3. Characterisation of yeast cells showing mitochondrial localisation.** Individual cells were segmented *and* isolated from confocal images for each strain and condition. The above images show recombined montages of 100 randomly selected cells from each individual cell image data set.

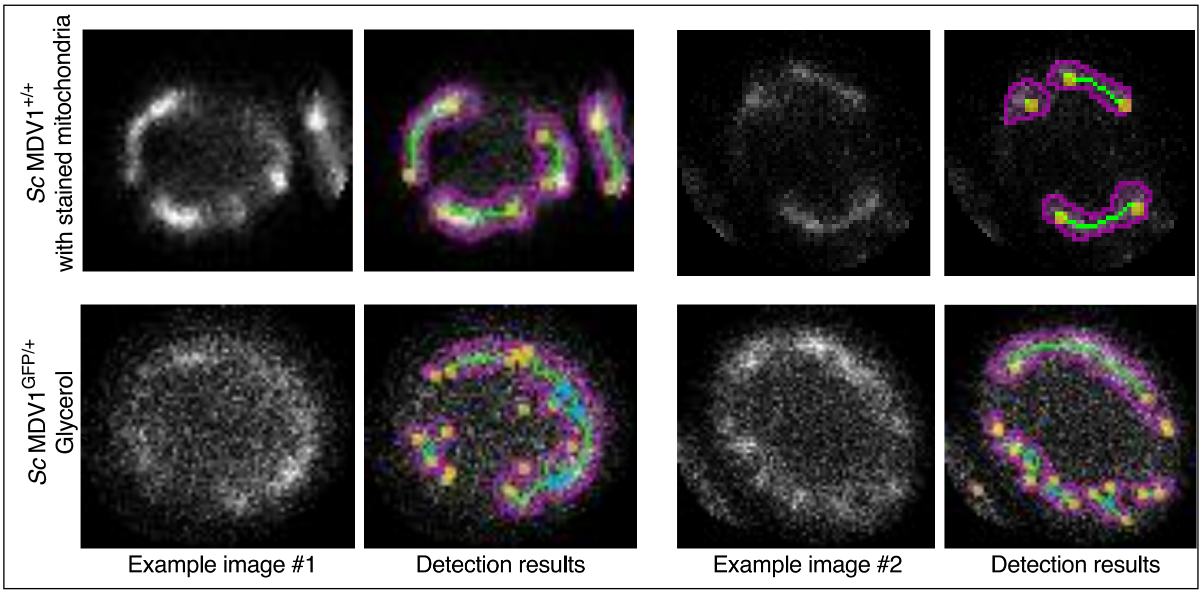


**Supplementary Figure 4. Quantification of cell size and mitochondrial surface in yeast.** Confocal microscopy images showing: Top panel; two *Sc* MDV1$^{+/+}$ cells with stained mitochondria. Bottom panel; two *Sc* MDV1$^{GFP/+}$ cells with stained mitochondria. Coloured areas were used to quantify the size of the cell and the localisation of mitochondria in the cells. *Sc* indicates *Saccharomyces cerevisiae*. Object detection was performed using ImageJ plugin Squassh (parameters: rolling ball window = 10, regularization = 0.05, minimum intensity = 0.15, noise model = poisson and objects below 2 pixels removed).

| Strain name used for FCS | Growth condition | Positive cells | Negative cells |
|---|---|---|---|
| *Sc* FIS1$^{GFP/+}$ | YP + 2 % Glucose | 682 | 1646 |
| *Sc* FIS1$^{GFP/+}$ | YP + 2 % Glycerol | 363 | 288 |
| *Sc* FIS1$^{GFP/-}$ | YP + 2 % Glucose | 783 | 1265 |
| *Sc* FIS1$^{GFP/-}$ | YP + 2 % Glycerol | 705 | 136 |
| *Sc* MDV1$^{GFP/+}$ | YP + 2 % Glucose | 436 | 187 |
| *Sc* MDV1$^{GFP/+}$ | YP + 2 % Glycerol | 661 | 50 |
| *Sc* MDV1$^{GFP/-}$ | YP + 2 % Glucose | 440 | 198 |
| *Sc* MDV1$^{GFP/-}$ | YP + 2 % Glycerol | 1042 | 68 |

**Table S2. Quantification of yeast cells showing mitochondrial localisation.** FCS: Fluorescence Correlation Spectroscopy; *Sc*: *Saccharomyces cerevisiae*; GFP: Green Fluorescence Protein; YP: yeast extract peptone medium
